# Poor sensitivity of iPSC-derived neural progenitors and glutamatergic neurons to SARS-CoV-2

**DOI:** 10.1101/2022.07.25.501370

**Authors:** Marija Zivaljic, Mathieu Hubert, Ludivine Grzelak, Giulia Sansone, Uwe Maskos, Olivier Schwartz

## Abstract

COVID-19 is a respiratory disease affecting multiple organs including the central nervous system (CNS), with a characteristic loss of smell and taste. Although frequently reported, the neurological symptoms remain enigmatic. There is no consensus on the extent of CNS infection. Here, we derived human induced pluripotent stem cells (hiPSC) into neural progenitor cells (NPCs) and cortical excitatory neurons to study their permissiveness to SARS-CoV-2 infection. Flow cytometry and western blot analysis indicated that NPCs and neurons do not express detectable levels of the SARS-CoV-2 receptor ACE2. We thus generated cells expressing ACE2 by lentiviral transduction to analyze in a controlled manner the properties of SARS-CoV-2 infection relative to ACE2 expression. Sensitivity of parental and ACE2 expressing cells was assessed with GFP- or luciferase-carrying pseudoviruses and with authentic SARS-CoV-2 Wuhan, D614G, Alpha or Delta variants. SARS-CoV-2 replication was assessed by microscopy, RT-qPCR and infectivity assays. Pseudoviruses infected only cells overexpressing ACE2. Neurons and NPCs were unable to efficiently replicate SARS-CoV-2, whereas ACE2 overexpressing neurons were highly sensitive to productive infection. Altogether, our results indicate that primary NPCs and cortical neurons remain poorly permissive to SARS-CoV-2 across the variants’ spectrum, in the absence of ACE2 expression.

## Introduction

As of April 2022, the COVID-19 pandemic had more than 453 million cases reported, and claimed more than 6 million lives. The number of deaths is largely under-estimated and may be closer to 18 million (1, 2).

COVID-19 was initially considered a respiratory disease. It soon became apparent that SARS-CoV-2 affects multiple organs including brain, and that neurological complications contribute to the disease’s acute and post-infectious symptomatology (3, 4). The potential of other respiratory viruses, including human coronaviruses (HCoVs): SARS-CoV-1, 229E, and OC43, and influenza virus, to cause neurological diseases were previously described (5-9). With SARS-CoV-2 infections increasing rapidly, the neurological symptoms were frequent and debilitating enough to attract a wide-spread attention of both the scientific community (3, 10) and general public, moving neurologists to the frontlines (Pleasure et al., 2020). Clinically distinct anosmia and ageusia were described in many cohorts (11). Clinical and pathological studies detected the presence of SARS-CoV-2 focally in patients’ brains in isolated cases (12-14). Further autopsies revealing viral presence in the CNS focused on the cortex (15). The neurotropic potential of SARS-CoV-2 was suggested with a human induced pluripotent stem cell (hiPSC) brain organoid model (15). Brain atrophy was reported in a large cohort of COVID-19 patients, as assessed by functional magnetic resonance imaging (fMRI) even in milder cases (16). Given the alterations in mental status (confusion, disorientation, agitation, and somnolence), collectively defined as encephalopathy, the unusual frequency of ischemic strokes, hemorrhages, seizures, the nervous system involvement became a critical aspect of COVID-19 (3, 17-20).

SARS-CoV-2 variants emerged in 2020, spreading at an increasingly rapid rate, exhibiting enhanced affinity of their spike towards the key cell entry receptor, angiotensin converting enzyme 2 (ACE2) (21), immune evasion and higher viral replication (22-24).

One of the key issues is the expression of ACE2 in neuronal cell types. RNA-seq analyses of human cortical brain tissue (25) (https://celltypes.brain-map.org) found very little evidence of *ACE2* expression across cell types in the brain. Another study (26) detected a limited to moderate *ACE2* expression in various brain regions. A low expression of *TMPRSS2 and furin*, key proteases of the infectious process, was found in neurons. *Neuropilin-1* (*NRP1*) RNA was detected in endothelial cells and developing brain, but mainly in adipose tissue and placenta (https://www.proteinatlas.org/). One study indicated NRP1 could significantly potentiate SARS-CoV-2 infectivity, finding it in infected cells of the nasal cavity, serving as a co-receptor (27). Of potentially greater clinical relevance is ACE2 expression in endothelial cells and pericytes. Their productive infection was described in a hiPSC model system (28). Particularly high levels of *ACE2* expression were found in the human choroid plexus (26). The productive infection and high incidence of syncytia reported in choroid plexus (29, 30) is likely to be responsible for viral entry and potential dissemination within the central nervous system (CNS).

However, different hypothesis of SARS-CoV-2 brain entry routes were proposed. It was suggested that in the nasal epithelium, infection of olfactory neurons extending axons towards the olfactory bulb could propagate the infection in the CNS. Transcellular transmission was described for other coronaviruses, specifically OC43 in a mouse model (31). An autopsy study found SARS-CoV-2 in the olfactory mucosa and hypothesized the axonal route of transport being a dominant one (32). This was not confirmed in subsequent studies (33). ACE2 and TMPRSS2 were detected in the nasal mucosa at both RNA and protein levels on sustentacular cells but not olfactory neurons (34). An early viremia related to COVID-19 was described (35, 36). The virus could thus potentially access the brain by crossing the blood-brain barrier. Viral RNA was found in the carotid artery of an individual during the acute phase of COVID-19 and the s protein was detected in cerebral and leptomeningeal endothelial cells (32), further supporting viremia and hematogenous route of transport of viral particles into the brain, where the blood-brain barrier is not intact.

We examined here SARS-CoV-2 entry and replication, on relevant cell types, hiPSC-derived cortical glutamatergic neurons and Neural Progenitor Cells (NPCs). The role of ACE2 to initiate a productive infection in neuronal cells infected with Wuhan, Alpha and Delta variants was tested, establishing the difference between viral susceptibility and a productive infection in neural cell types.

## Methods

### HiPSC culture

Human induced pluripotent stem cell (hiPSC) lines are from the Wellcome Trust Sanger Centre collection: (https://www.sanger.ac.uk/collaboration/hipsci/, line WTS2). The methods to derive these cells have been described previously (37). Briefly, hiPSC colonies are maintained in Essential 8™/E8 Medium on 6-well plates coated with Geltrex. The hiPSC colonies were detached by treatment with ReLeSR (Stemcell Technologies) and kept in suspension in low-attachment plates (Corning) as embryoid bodies (EBs) during 5 days in EB media consisting of Advanced DMEM-F12, KnockOut™ Serum Replacement, Penicillin-Streptomycin, MEM Non-Essential Amino Acids Solution, 2-Mercaptoethanol, GlutaMAX™, B27 without vitamin A and N2 supplements, additionally supplemented with 10 μM SB421542 (Sigma-Aldrich) and 20 μM dorsomorphin (Sigma-Aldrich) to induce the formation of neuroectoderm. EBs are then plated in dishes coated with polyornithine (Sigma Aldrich) and laminin (Sigma Aldrich) and kept in culture for 7 days in EB media until neural rosettes were observed. These rosettes were then picked manually and plated in 60×15 mm PLO/lam-coated petri dishes and kept in culture for 7 days in neural precursor media (Advanced DMEM-F12, Penicillin-Streptomycin, 2-Mercaptoethanol, B27 minus Vitamin A, N2 and GlutaMAX™ supplements and growth factors FGF2 [20 ng/μl], and EGF [20 ng/μl]). On the day 21, the neural rosettes were picked manually and dissociated into neural progenitor single-cell suspension using StemPro™ Accutase™ Cell Dissociation Reagent. They are then plated in PLO/lam-coated 96-well-plates at a density of 1.5 × 105 cells/mm2 and kept in neuronal differentiation media for 5 weeks (Neurobasal medium, B27 without vitamin A, N2 and GlutaMAX™ and CultureOne supplements, Penicilin-Streptomycin, dibutyryl cAMP [100 mM], L-Ascorbic acid [0.2 M], laminin (1 mg/ml) and BDNF [10 ng/μl], GDNF [10 ng/μl]. Half of the used media volume was substituted with fresh media twice a week. Products were from Thermo Fisher unless stated otherwise. The cultures were regularly tested for mycoplasma.

### Generation of ACE2 stably expressing cells

For transduction with the lentiviral vector expressing ACE2, 2 × 10^4^ cells were resuspended in 150 µl of NPC media containing 5–25 μl of lentiviral vectors. Cells were agitated 30 s every 5 min for 2.5 hours at 37°C in a Thermomixer. Media was then topped to 1.5 ml. The cells were plated in PLO/lam-coated 6-well plates and incubated for 48h at 37°C.

### Production of lentiviruses and pseudo-infection assays

Pseudotyped non-replicative lentiviral vectors were a kind gift from Dr. Charneau’s laboratory at the Institut Pasteur. They were produced by transfection of 293T cells (38). Briefly, cells were co-transfected with plasmids encoding for lentiviral proteins; a GFP or a luciferase reporter and the SARS-CoV-2 plasmid encoding for spike protein or the VSV-g plasmid to be used as a control. Lentiviral vector to deliver the gene encoding ACE2 under the transcriptional control of the EF1α was previously described (39). Supernatants with pseudotyped virions were harvested at 48 h post-transfection, clarified by 6-min centrifugation at 2500 rpm at 4°C. Production efficacy was assessed by measuring infectivity or p24 concentration.

For infectivity assays, different concentrations of lentiviruses were added to the media of the cells cultured in 96-well plates (15000 cells per well) and incubated for 48h. Half of the virus containing media was aspirated and replaced with luciferase substrate (Bright-glo luciferase assay system, Promega) and incubated for 5 minutes at 37°C. Bioluminescence emission signal was then acquired with the Perkin Elmer spectrometer. Cultures infected with GFP-labelled pseudotyped lentiviruses were fixed after 48h and imaged on Opera Phenix confocal microscope.

### Virus

Experiments with SARS-CoV-2 isolates were performed in a BSL-3 laboratory, following safety and security protocols approved by the risk prevention service of Institut Pasteur. The Wuhan SARS-CoV-2 strain (BetaCoV/France/IDF0372/2020) and the D614G strain (hCoV-19/France/GE1973/2020) were supplied by Dr. S. van der Werf of the National Reference Centre for Respiratory Viruses (Institut Pasteur, Paris, France). The D614G viral strain was sourced through the European Virus Archive goes Global (EVAg) platform, which is funded by the European Union’s Horizon 2020 research and innovation program under grant agreement 653316. The Alpha strain was isolated in Tours, France, from an individual who returned from the United Kingdom (40). The Delta strain was isolated from a nasopharyngeal swab of a hospitalized patient returning from India (41). The swab was provided and sequenced by the Laboratoire de Virologie of the Hopital Européen Georges Pompidou (Assistance Publique des Hôpitaux de Paris). Informed consent was provided by the individuals for use of their biological materials. The viruses were isolated and amplified by one or two passages on Vero cells. The viruses were sequenced directly from the nasal swabs and upon passaging. Titration of viral stocks was performed by 50% tissue culture infectious dose (TCID50).

### Flow cytometry

Surface staining of ACE2 in NPCs and cortical neurons was performed before fixation, in PBS 1% BSA. Cells were incubated with primary ACE2 antibody (AF933—R&D) for 30 minutes at 4°C, washed twice with PBS and then incubated with secondary Alexa Fluor 647 antibody (Invitrogen) in 1:500 dilution for 30 min at 4°C. Cells were fixed for 15 min in 4% PFA and acquired on an Attune NxT Flow Cytometer (Life Technologies). The data was analyzed using FlowJo software (Tree Star, OR, USA).

### SARS-CoV-2 inoculation, immunocytochemistry and microscopy

Cells were infected with different MOI of SARS-CoV-2 variants in 50 µl of media and incubated for 2 hours. The viral inoculum was then removed, cells were washed twice with PBS and 100 µl of media was added. At 2, 24, 48, 72 and 144 hours post-infection, cells were fixed in 4% PFA for 30 min at room temperature and washed in PBS.

Cellular and spike (S) proteins were labelled with specific antibodies. Cells were stained with the indicated primary antibodies for 45 min in PBS, 1% BSA, 0.05% sodium azyde, and 0.05% saponin. Cells were then washed twice with PBS, 0.05% saponin and stained with secondary antibody for 30 min at RT in PBS, 1% BSA, 0.05% sodium azyde, 0.05% saponin, and Hoechst 33342 (1:10,000) and washed with PBS. Microscopy images were acquired on Opera Phenix microscope (PerkinElmer) and analyzed on Harmony High-Content Imaging and Analysis Software.

The following antibodies were used: Anti-spike antibody mAb102 (Planas et al., 2021a); nestin (SP103 Invitrogen), MAP2 (ab5392), β-III tubulin (ab7751) and VGlut1 (Synaptic Systems) according to the manufacturer’s instructions. Secondary antibodies coupled with Alexa Fluor 488 or 647 (Invitrogen) were applied at 1:500 for ICC.

### Western blot

Cells were harvested (10-15 million) and lysed in TXNE buffer (1% Triton X-100, 50 mM Tris– HCl (pH 7.4), 150 mM NaCl, 5 mM EDTA, protease inhibitors) for 30 min on ice. Equal amounts (50 μg) of cell lysates were analyzed by western blot. The following antibodies were diluted in wb-buffer (PBS, 1% BSA, 0.05% Tween, 0.01% Na azyde) goat anti-human ACE2 (R&D AF933, 1:2,000), rabbit anti-human TMPRSS2 (Atlas antibodies HPA035787, 1:1,000), rabbit anti-human actin (Sigma A2066, 1:2,000). Specie-specific secondary DyLight-coupled antibodies were used (diluted 1:10,000 in wb-buffer) and proteins were revealed using a Licor Imager. Images were processed using Image Studio Lite software.

### Viral Kinetics and Release

Viral RNA in supernatants of infected cells and intracellular RNA were extracted using Quick-RNA™ Viral 96 Kit (Zymo Research). RT-qPCR was performed from 1µL of template RNA in a final volume of 5 μL per reaction in 384-well plates using the Luna Universal Probe One-Step RT-qPCR Kit (New England Biolabs) on a QuantStudio 6 Flex thermocycler (Applied Biosystems). Reactions were performed in triplicates, using SARS-CoV-2 nucleocapsid (n) protein specific primers and human ribosomal protein L13 (hRLP13) as an endogenous reference control gene.

The differences in the CT SARS-CoV-2 n amplicons and the CT of the endogenous reference control, representing ΔCT, were calculated to normalize the quantity of nucleic acids. The fold change in gene expression was calculated using 2^-ΔCT^.

Viral RNA in the extracellular compartment was assessed and quantified using standard curve which was set up in parallel with purified SARS-CoV-2 viral RNA.

Infectious virus release was assessed by harvesting supernatant at each time point and performing a TCID50 assay with Vero cells.

### Statistical analyses

Flow cytometry data was analyzed with FlowJo v10 software (Tree Star). Calculations were all performed with Microsoft Excel 365. GraphPad Prism 9 was used to generate figures and for statistical analysis. Statistically significant difference between multiple conditions was evaluated using nonparametric Kruskal-Wallis test.

## Results

### Generation and characterization of HiPSC derived NPCs and cortical neurons

HiPSCs (WTS2 line, Wellcome Trust) were derived through the formation of embryoid bodies (EBs) first into NPCs and then into cortical glutamatergic neurons (Fig. 1A). Immunocytochemical (ICC) labelling for NPCs and cortical glutamatergic neurons’ specific antigens detected nestin in NPC cultures and MAP2, β-III tubulin and VGlut1 in cultures of glutamatergic neurons (Fig. 1B). We assessed ACE2 expression patterns in NPC and neuronal cultures by flow cytometry and Western Blot. We did not detect ACE2 in wild-type (WT) cells, in contrast to the NPC cell line stably transduced with a lentiviral vector coding for ACE2 (Fig. 1C, D). The ACE2+ NPCs were in parallel differentiated into cortical glutamatergic neurons, which retained ACE2 expression (Fig. 1C, D).

**Figure 1.**
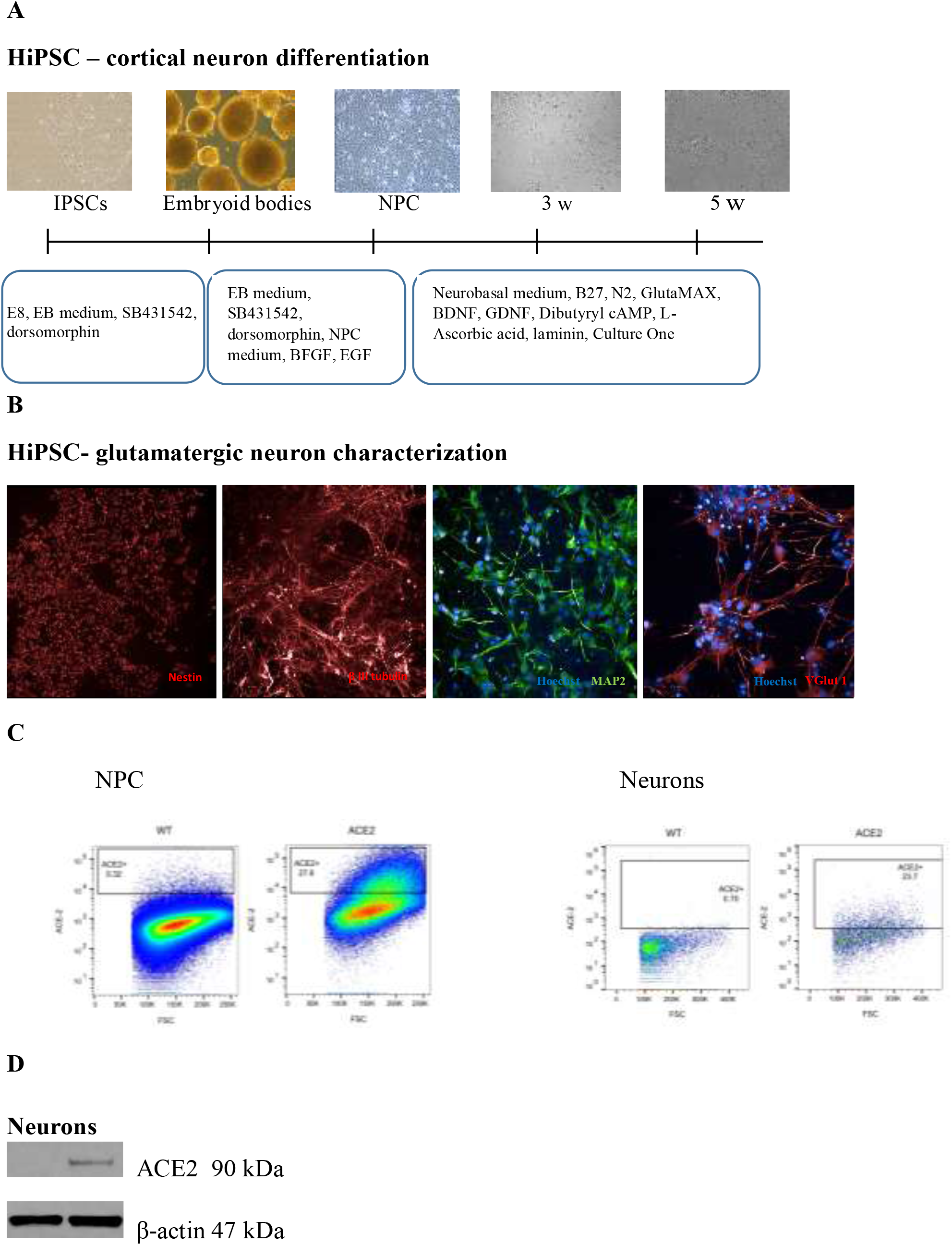
Generation and characterization of NPCs and cortical glutamatergic neurons. HiPSC-neuron differentiation, morphological characterization and ACE2 expression of hiPSC-derived neuronal progenitor cells (NPCs) and cortical neurons. (A) HiPSC were derived into NPCs through the formation of embryoid bodies. NPCs were further differentiated toward the cortical fate. An immunofluorescence staining was performed after differentiation. (B) NPCs were labelled with nestin and cortical excitatory neurons were MAP2, beta III-tubulin and VGlut1 positive. Images were acquired on Opera Phenix confocal microscope (40x). Scale bars: 10-50 µm. (C) Flow cytometry analysis of ACE2 levels in endogenously expressing (termed WT here) and ACE2 transduced NPCs and neurons. (D) ACE2 expression in WT and ACE2 transduced neurons assessed by western blot; β-actin was used as a control protein. Results are representative of at least two independent experiments.

### S-pseudotyped lentiviral infections of NPCs and cortical neurons assessing S-mediated cell entry

We first tested the sensitivity of NPCs and cortical neurons in ACE2 negative and positive conditions using S-pseudotyped single-cycle lentiviruses (termed HIV (spike) vectors) with GFP or luciferase reporters (Fig. 1A). HIV (VSV) vectors were used as a positive control of infection. GFP-encoding vectors were tested in order to visualize infection by microscopy (S1). Assays with luciferase-encoding vectors were then used, offering higher detection sensitivity. WT NPCs and neurons, not expressing ACE2, were not sensitive to infection (Fig. 2A, B and Fig. S1). ACE2 expressing cells were infected, with a correlation between the viral inoculum and detected signal (Fig. 2A, B). ACE2 transduced NPCs were about 6-times more sensitive to infection than cortical neurons differentiated from the same batch. The two cell types expressed similar levels of ACE2 (Fig. 1C, D). The cells were infected with HIV (VSV) in a parallel, control experiment to test cellular viability and sensitivity to infection (Fig. 2C, D). HIV (VSV) infected cells irrespectively of the presence of ACE2, and was about 2000 times more infectious than S-pseudotyped lentiviruses in the ACE2+ condition, as detected by luciferase production.

**Figure 2.**
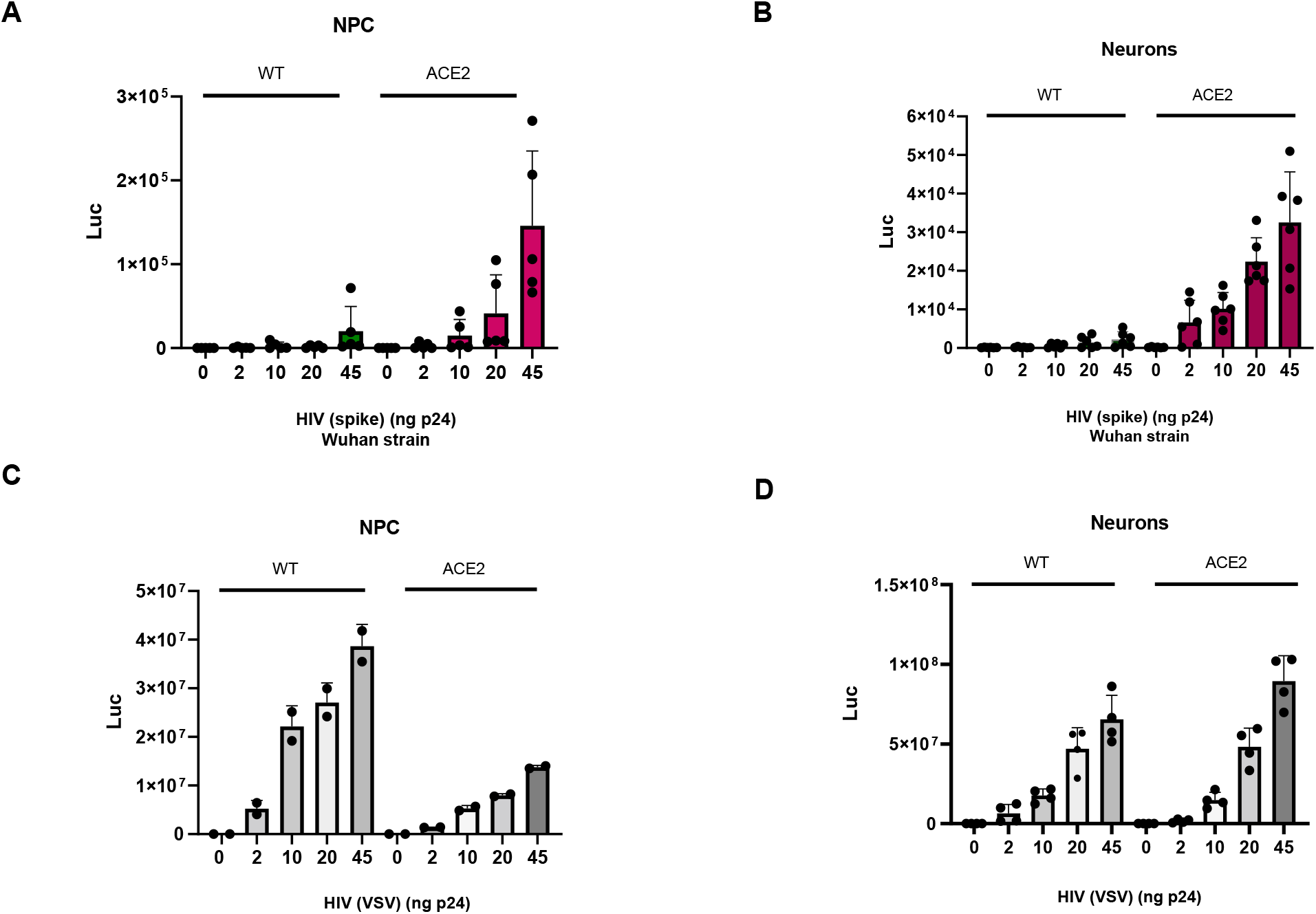
Sensitivity of WT and ACE2-expressing NPCs and neurons to spike-carrying pseudovirus. S-pseudotyped lentiviral infections of NPCs and neurons and HIV (VSV) control infections. Bioluminiscence assays were performed, luciferase emission measured 48h after infection and represented on the graphs. A) WT and ACE2 transduced NPCs were infected with s-pseudotyped Wuhan variant lentivirus, B) WT and ACE2 transduced neurons were infected with the same virus. The data represents 2 independent experiments conducted in triplicates. C) WT and ACE2 transduced NPCs were infected with HIV (VSV) virus as a control experiment conducted in duplicates, D) WT and ACE2 transduced neurons were infected with the same virus in 2 control experiments conducted in duplicates. Titration was performed using the p24 assay. 10^−3^ 10^−2^ 10^−1^

We then examined the sensitivity of NPCs to infection by pseudoviruses carrying Wuhan, D614G, Alpha or Beta spike proteins (Fig. S2), given the increased affinity of Alpha and Beta for ACE2, when compared to ancestral strains (Ramanathan et al., 2021). In WT NPCs, luciferase emission seemingly varied but overall gave very low values across the pseudotyped variants tested. The luciferase signal remained 500-2000 times lower than measured with HIV (VSV) (Fig. 2C and Fig. S2). The addition of ACE2 in NPCs renders the cells sensitive to pseudotypes carrying different variants’ spikes. There were no statistically significant differences in the infectivity rates between the variants, in the absence or presence of exogenous ACE2 (Fig. S2A, B).

### Sensitivity of NPCs and neurons to infectious SARS-CoV-2 ancestral strains

Whether NPCs allow productive infection with authentic SARS-CoV-2 was examined next. We first used the ancestral D614G strain. WT and ACE2+ NPCs were exposed to a range of MOIs. Cells were stained one day post-infection, with an anti-spike antibody (mAb102), counter-stained with Hoechst and imaged by confocal microscopy (Fig. 3A). The ACE2+ NPCs were readily infected and formed syncytia which increased in number with the multiplicity of infection (MOI). On the contrary, WT NPCs were not sensitive to infection (Fig. 3A). Similar results were obtained at later time points.

**Figure 3.**
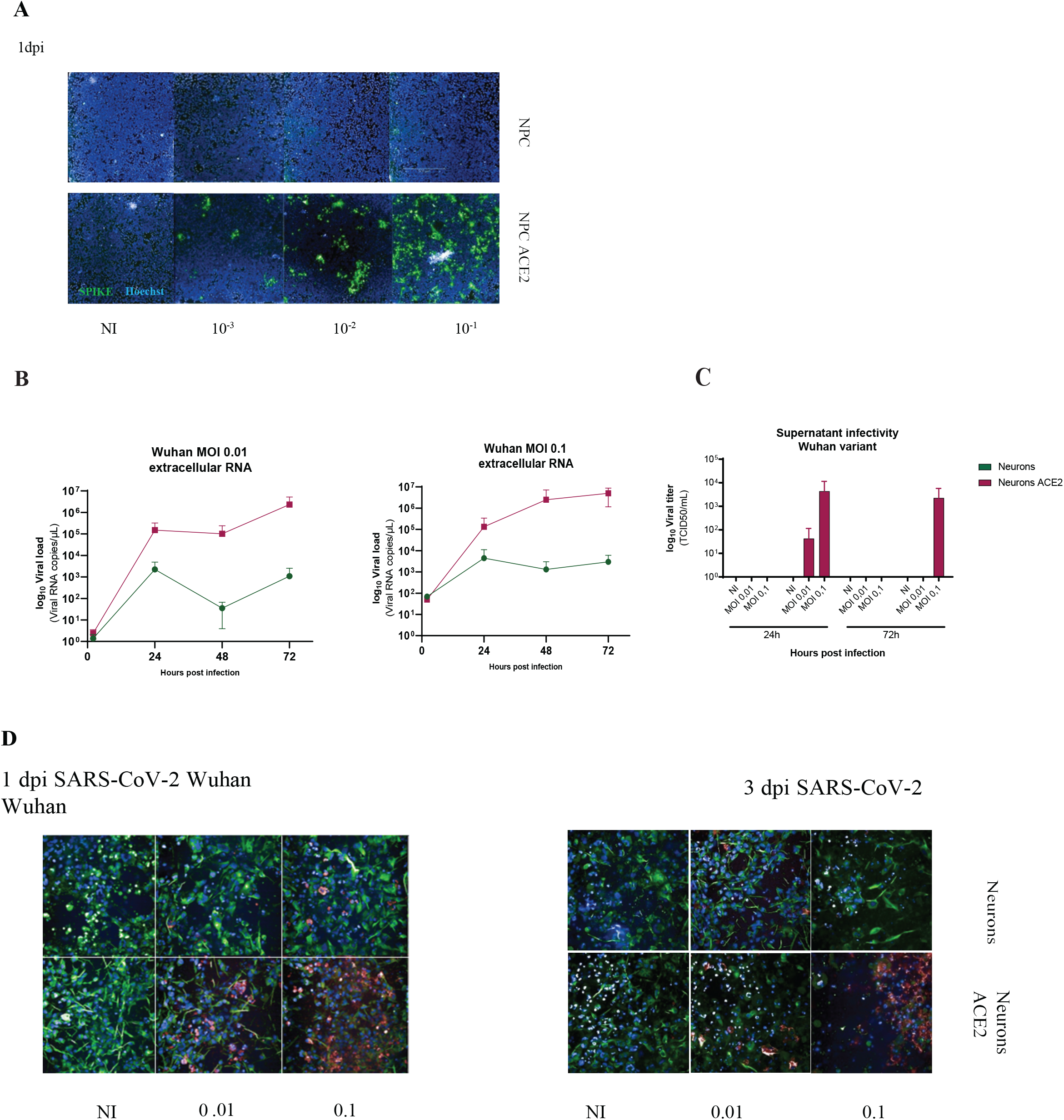
Sensitivity of NPCs and neurons to infectious SARS-CoV-2. (A) Immunofluorescence analysis of ACE2-expressing NPCs infected with SARS-CoV-2 (ancestral D614G strain). Samples were fixed 1 day after infection and immunostained for spike. Data are representative of 6 independent experiments (B). Kinetic analysis of viral RNA release by endogenously expressing WT and ACE2-expressing cortical neurons. Cells were infected with SARS-CoV-2 (ancestral D614G strain) at two multiplicities of infection (MOI). Supernatants were harvested at the indicated days post infection. Levels of SARS-CoV-2 RNA were measured by RT-qPCR. Data are mean ±SD of triplicates from 2 independent experiments. (C) Release of infectious virus in the supernatants: WT and ACE2-expressing cortical glutamatergic neurons were infected with SARS-CoV-2 ancestral Wuhan strain at two multiplicities of infection (MOI). Supernatants were harvested at the indicated days post infection. Levels of infectious virus were measured in a plaque assay on Vero E6 cells. Data are mean ±SD of triplicates from at least 2 independent experiments. (D) Immunofluorescence analysis of WT and ACE2-expressing glutamatergic neurons infected with SARS-CoV-2 ancestral Wuhan strain. Cells were fixed 1, 2, 3 and 6 days post infection, immunostained with MAP2, spike antibodies and Hoechst. The images were taken on the Opera Phenix confocal microscope (40 x). Data are representative of 3 independent experiments.

We then differentiated NPCs toward cortical glutamatergic neurons. Following differentiation, WT and ACE2+ cortical neurons were inoculated with SARS-CoV-2 Wuhan strain at two MOIs. Viral release in the supernatant was assessed by RT-qPCR at different time-points post infection (Fig. 3B). Viral RNA in the supernatant of ACE2+neurons increased up to 10^5^-fold one day post infection (pi) and remained high at days 2 and 3 pi. Low levels of viral RNA were detected in the supernatants of WT neurons (Fig. 3B). We measured release of infectious viral particles in supernatants of infected cells by a TCID50 plaque assay at days 1 and 3 pi (Fig. 3C). Infectious virus was detected in ACE2+ neurons, and not in WT cells. Infectivity in the supernatants appeared to be higher at day 1 than day 3 pi (Fig. 3C). We also assessed the morphological aspect of neurons upon infection. The cells were fixed at different time-points and labelled with spike and neuron-specific antibodies (Fig. 3D). WT neurons were sporadically spike positive with no evidence of progressive infection. ACE2+ neurons were vastly infected at both MOIs. A drastic decrease in cell viability was observed at the later time-point in ACE2+ cells and not in WT cells (Fig. 3D).

Altogether, these results indicate that WT NPCs and glutamatergic neurons are not sensitive to efficient productive SARS-CoV-2 infection with ancestral Wuhan and D614G strains. The low levels of viral RNA detected in the supernatants did not correspond to infectious viral particles. ACE2+ neurons are sensitive to infection, release high levels of the viral RNA and infectious viral particles, and die upon infection.

### Sensitivity of neurons to Alpha and Delta SARS-CoV-2 variants

The Alpha and Delta SARS-CoV-2 variants had replaced previous strains worldwide during 2020 and 2021. They display increased affinity towards ACE2 and infectivity (24, 41-43). We thus compared sensitivity of WT and ACE2+ neurons to infection by Alpha and Delta variants (Fig. 4). The Wuhan strain was included as a control. The three strains were inoculated at two MOIs. RNA was extracted and viral replication was assessed by measuring intracellular viral RNA at 4 time points (2, 24, 48 and 72h pi) and by confocal microscopy. In ACE2+ neurons, upon infection with the three viral strains, efficient viral replication and spike-expressing cells were observed (Fig. 4). In contrast, none of the viral isolates efficiently replicated in WT neurons (Fig. 4). Low levels of viral RNA and only sporadic spike-positive cells were detected by confocal microscopy after exposure to Alpha and Delta variants (Fig. 4B), as observed when neurons were exposed to the ancestral Wuhan strain (Fig. 3D). Their typical morphology remained intact, neurons sporadically infected with Alpha and Delta variants were clustered, as compared to the ones infected with Wuhan variant (Fig. 4A). With the three variants, infection was widespread and destructive to the neuronal networks formed when ACE2 receptor was expressed (Fig. 3, 4A). High incidence of multinucleated syncytia was evident upon infection with Alpha and Delta variants, due to their higher affinity towards ACE2 receptor (Fig 4A). S immunostaining showed high intensity in ACE2 neurons due to extensive viral replication and accumulation of viral particles, in line with viral RNA levels. The variants replicated at low levels in the WT condition (Fig. 4B, C, D), with Wuhan and Alpha variants reaching a peak at 24 h (Fig. 4B, C) and Delta variant at 72 h (Fig. 4D). There was 10^2^-10^3^-fold increase in viral replication in ACE2 over-expressing neurons, in comparison to the WT condition, with the three variants. Therefore, neurons do not allow efficient viral replication and productive infection, without addition of exogenous ACE2.

**Figure 4.**
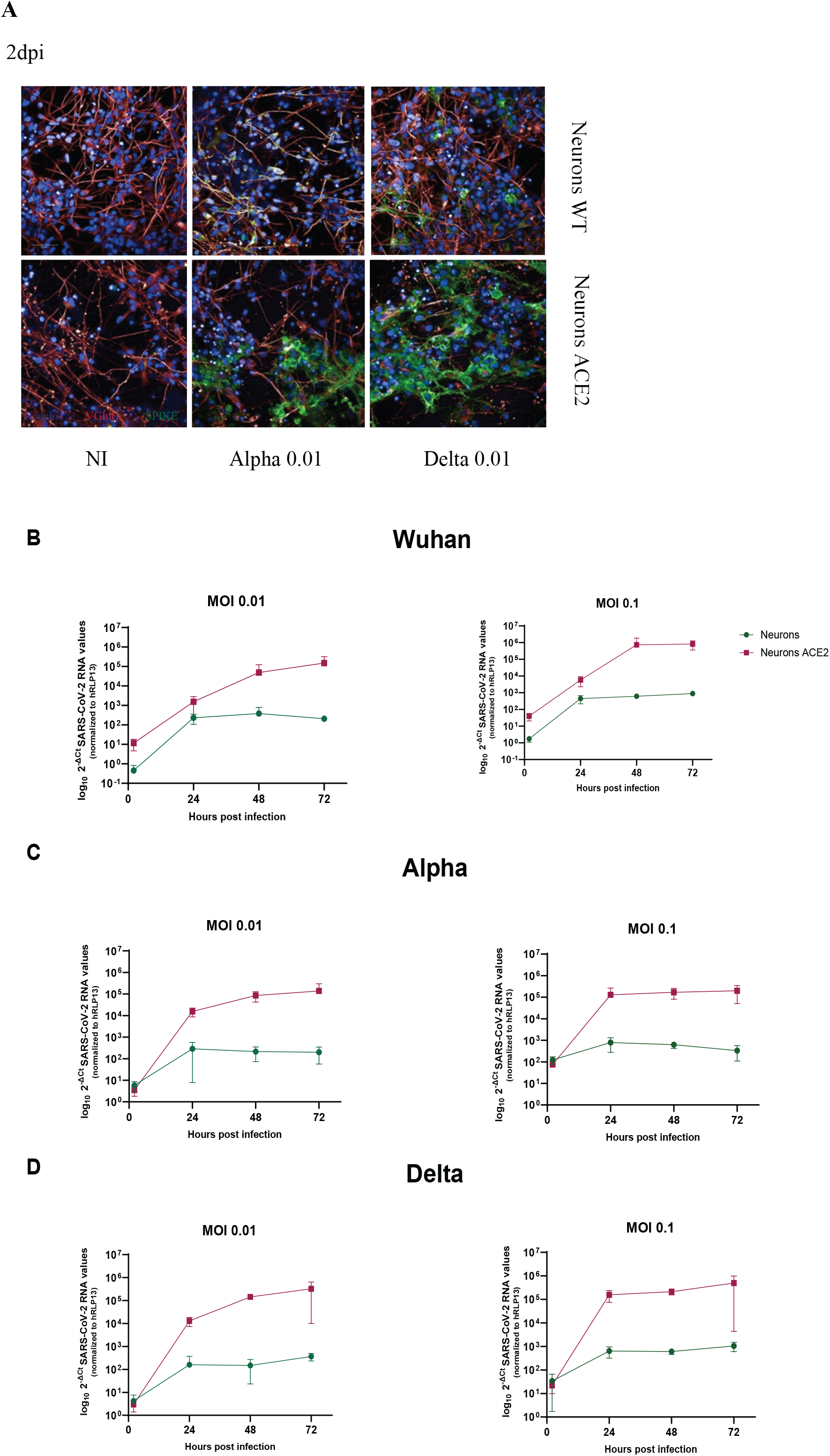
Sensitivity of WT and ACE2-expressing neurons to SARS-CoV-2 Alpha and Delta variants. (A) Immunofluorescence analysis of WT and ACE2-expressing neurons infected with SARS-CoV-2 ancestral Wuhan strain. Cells were fixed 1, 2 and 3 days post infection, labelled with VGlut1 and DAPI, spike antibodies and Hoechst. The images were taken on the Opera Phenix confocal microscope (40 x). Data is representative of 2 independent experiments in triplicates. (B) Kinetic analysis of viral RNA release by WT and ACE2-expressing cortical neurons. Cells were infected with SARS-CoV-2 Alpha and Delta variants at two MOI. Supernatants were harvested at the indicated days post infection. Levels of SARS-CoV-2 RNA were measured by RT-qPCR. Data are mean ±SD of triplicates from 5 independent experiments.

## Discussion

The pathophysiological mechanisms of neurological COVID-19 manifestations, SARS-CoV-2 neurotropism and its possible consequences are not well understood. Immortalized cell lines and animal models have been used in SARS-CoV-2 research, but they are relatively remote from human biology and are less likely to capture all features of viral infection than primary cells. Most of human cell lines used in vitro carry mutations in proteins associated to tumors, such as P53, that could regulate SARS-CoV replication (44). This raises concerns about the use of immortalized cancer cell lines in SARS-CoV-2 and fundamental research and drug development. HiPSC models have been used to study neurotropic viruses such as Zika virus and Herpes Simplex Virus 1 (HSV1) (45). HiPSC model systems were developed to accurately recapitulate host-viral interactions and study the sensitivity of neuronal cell types of SARS-CoV-2 infection.

We found that iPSC-derived NPCs and cortical glutamatergic neurons are not at all or poorly sensitive to SARS-CoV-2 infection. By transducing the cells with a vector encoding ACE2, we rendered NPCs and neurons able to replicate SARS-CoV-2 and release infectious virus. Similar results were obtained with Wuhan and D614G ancestral viral strains and variants Alpha and Delta. Our work confirms and extends previous results using iPSC derived neural cells reporting absence of productive SARS-CoV-2 infection (Jacob et al., 2020, Pellegrini et al., 2020).

Sensitivity of these primary cells to SARS-CoV-2 was explained further. The receptor ACE2 is likely a limiting step of neuro-invasion and its low expression prevents robust infection of brain parenchyma. We did not observe an increase of infection over time where ACE2 is expressed endogenously, suggesting that trans-cellular viral transport is not efficient in the absence of ACE2. No correlation between TMPRSS2 and NRP1 expression and infectivity rates in brain organoids has been reported (Song et al., 2020) suggesting that neurons cannot be infected through alternative entry pathways. Moreover, detection of viral RNA in human tissues does not necessary correspond to the presence of an active site of viral replication, as illustrated here by the presence of low viral RNA loads without infectious virus measured in TCID50 assays.

Glutamatergic neurons make a relevant model given that the disruption in glutamate homeostasis and excitotoxicity leading to degeneration and cell death was demonstrated in several viral infections, such as West Nile virus, Sindbis virus and Japanese Encephalitis virus (46-48), and in an animal model of HCoV-OC43 infection (49). Given that the SARS-CoV-2 cases of vertical transmission were reported, tropism of NPCs was tested (50-52). NPCs mimicked a developing telencephalon and may serve as a model system to explore potential effects of SARS-CoV-2 on the developing brain, when the blood-brain barrier is not yet well formed. SARS-CoV-2 infection of NPCs gave negative results, suggesting that the developing brain may not be a target for the virus.

However, even though hiPSC models provide an accurate perspective into infectious mechanisms and host-viral interactions, they do not give insights into systemic effects or indirect consequences of the infection, important to consider given the multifaceted character of COVID-19. SARS-CoV-2 may cause clinical signs of encephalitis (12, 13, 53-56) with defined criteria such as altered mental status, fever, seizures, white blood cells in the CSF and focal brain abnormalities by neuroimaging (57). Several autopsy series have revealed a notable lack of productive infection and immune cell infiltration, therefore showing no evidence of encephalitis (14, 58, 59). Possible artifact RNA contamination was raised, given the challenge of dissecting the brain parenchyma from vasculature and meninges (Solomon et al., 2020).

Although the lack of detection of immune cells surrounding infected cells excludes a classic viral encephalitis with vast microglial activation, the occurrence of sub-acute and sub-clinical infection cannot be excluded. A detailed post-mortem analysis showed robust antibody response in the CNS, localized ischemia and cell death (Solomon et al., 2020). Whether a minimal infection of neurons in humans may lead to cell death or viral clearance remains to be explored. One study identified dopaminergic neurons as a more susceptible target (60). Therefore, long COVID-19 association with brain infection has been suggested by analogy to other respiratory viruses (61), but so far evidence points that sequalae are probably mainly due to chronic sub-hypoxia, metabolic dysfunction, hormonal and auto-immune dysregulations (62).

Consistent with hypoxic brain injury, autopsy studies in COVID-19 have shown neuronal damage in regions most vulnerable to hypoxia, including neocortex, hippocampus, and cerebellum (Kantonen et al., 2020; Reichard et al., 2020; Solomon et al., 2020). These damages are in line with recent fMRI volumetric findings indicating atrophic changes within the same areas in COVID-19 patients, as compared to their pre-pandemic scans and contrasted to results of healthy individuals (Douaud et al., 2022). A dramatic decrease in transthyretin (TTR) production was reported (Jacob et al., 2020). TTR is a necessary transporter of the thyroid hormone thyroxine from blood to CSF. Moreover, lower TTR production by the choroid plexus has been described in neuropsychiatric disorders (63) leading to low concentrations of thyroxine in CSF, associated with neurological symptoms such as decreased attentiveness, slowing of thought and speech and deteriorated cognition, all reported in COVID-19 patients (Douaud et al., 2022,).

Another cause of brain alteration may be the local production of inflammatory cytokines, normally recruiting immune cells to infection sites and are an important piece in COVID-19 pathophysiology. Increased circulating levels of IL-6, IL-1β, and TNF-α, as well as IL-2, IL-8, IL-17 are present in most cases, and serum levels of IL-6 and TNF-α reflect disease severity (64). It could be worth measuring RNA levels of inflammatory cytokines in more elaborated assembloid-systems ensuing SARS-CoV-2 exposure.

Brain damage may also be linked to thrombosis. The risk for stroke in hospitalized COVID-19 patients is 7-fold higher than after influenza infection (Merkler et al., 2020). Severe SARS-CoV-2 infection is associated with high risk of hemagglutination, with elevated D-dimer and fibrinogen (Helms et al., 2020). Autopsy series demonstrated wide-spread micro-thrombi associated with patchy acute brain infarction, with hemophagocytosis and disproportional vascular congestion (65).

In conclusion, our results confirm that SARS-CoV-2 should not be considered as a virus primarily attacking the nervous system, such as HSV-1. We show that NPCs and neurons are equipped with all components allowing SARS-CoV-2 replication but lack sufficient expression of the critical receptor ACE2. In most cases, neurological signs associated with acute, severe, or long COVID-19 are unlikely due to a productive infection of neurons but are rather linked to indirect mechanisms.

## Acknowledgements

The authors would like to thank all the well-wishers. Members of Virus and Immunity Unit and Integrative Neurobiology of Cholinergic Systems Unit for support, Nathalie Aulner, Anne Danckaert and the UtechS Photonic Bioimaging (UPBI) core facility (Insitut Pasteur), a member of the France BioImaging network, for image acquisition and analysis support. Work in UPBI is funded by grant ANR-10-INSB-04-01 and Région Ile-de-France program DIM1-Health. Work in OS lab is funded by Institut Pasteur, Fondation pour la Recherche Médicale (FRM), Urgence COVID-19 Fundraising Campaign of Institut Pasteur, ANRS, the Vaccine Research Institute (ANR-10-LABX-77), Labex IBEID (ANR-10-LABX62-IBEID), ANR/FRM Flash Covid PROTEO-SARS-CoV-2 and IDISCOVR. Hugo Mouquet for the kind gift of antibodies, the TheraVectys team for providing all the lentiviral vectors used in this study and Vinita Jagannath for HiPSC culture training, Nicoletta Casartelli for titration. LG is supported by the French Ministry of Higher Education, Research and Innovation. MZ was a scholar of the Pasteur-Paris University (PPU) International Doctoral Program, and is supported by the Institut Pasteur Bourse and ERANET. UM’s laboratory is funded by the Urgence COVID-19 Fundraising Campaign of Institut Pasteur, ANR RA Covid-19, FRM EQUIPE 2019, and ERANET iPS&Brain. The funders of this study had no role in study design, data collection, analysis, interpretation, or the writing of the article.

## Author contributions

Experimental strategy and design: MZ, MH, LG, OS

Experimentation: MZ, MH, LG, GS

Vital materials, expert advice and training: PC, UM, VJ

Data processing and figure generation: MZ, MH, LG

Manuscript writing and editing: MZ, OS

Supervision: OS

## Conflict of interests

The authors have no conflict of interest.

## Supplementary figures

**Figure S1.**
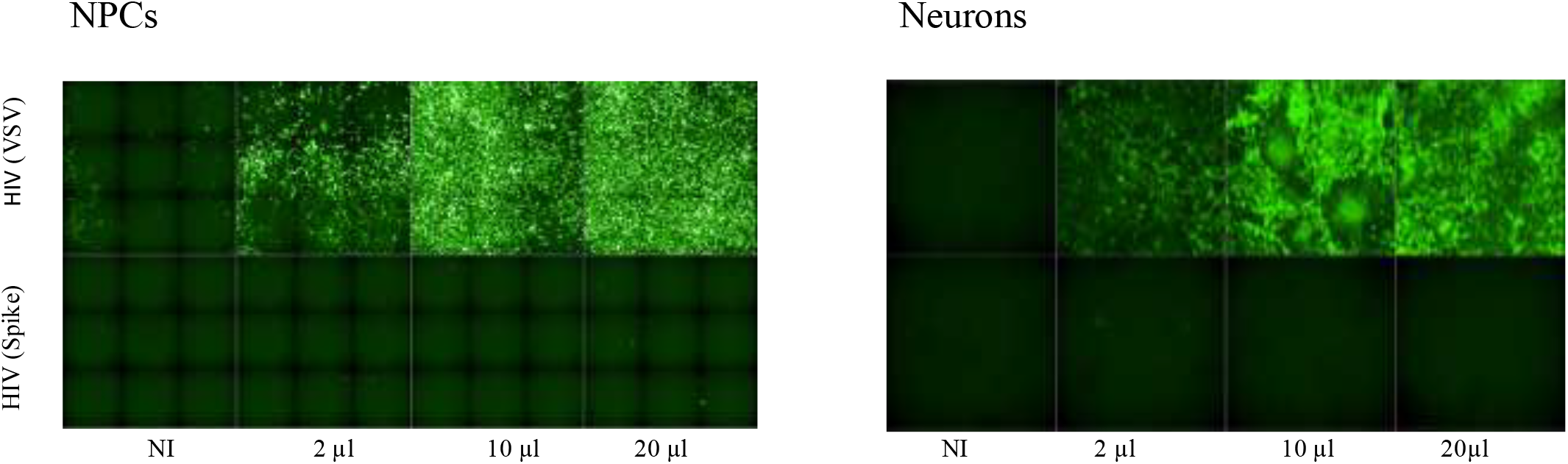
NPCs and neurons are not sensitive to GFP-expressing spike-carrying pseudovirus. WT NPCs were infected with the indicated amounts HIV (spike) pseudotypes carrying the Wuhan spike. HIV (VSV) was used as a control. The cells were fixed 48h after infection and imaged on Opera Phenix confocal microscope (10x) for visualization. Cells are sensitive to HIV (VSV) but not to HIV (spike) pseudotypes. Data are representative of 2 independent experiments.

**Figure S2.**
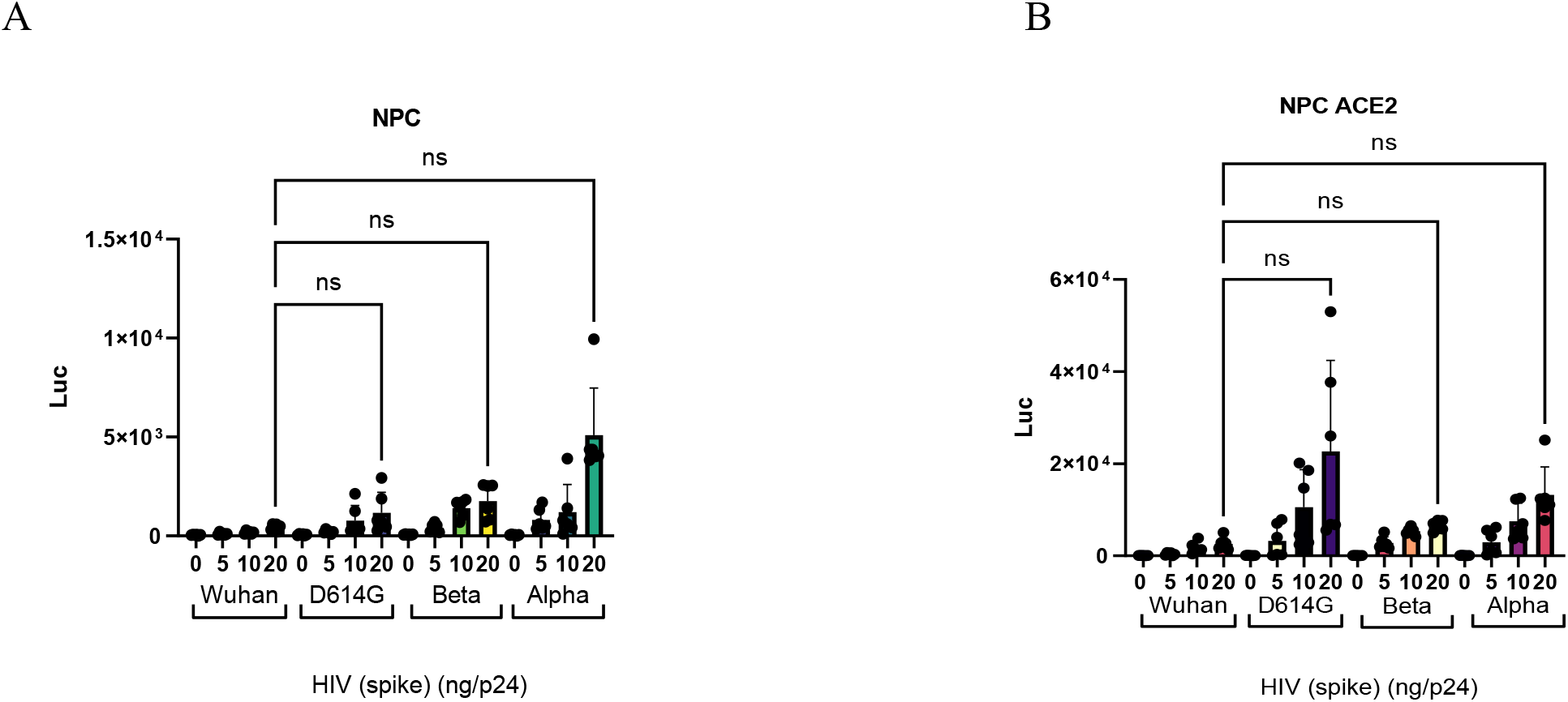
Sensitivity of WT and ACE2-expressing NPCs pseudovirus carrying SARS-CoV-2 variant spikes. WT and ACE2 transduced NPCs were infected with the indicated amounts of HIV (spike) pseudotypes (in ng of p24) carrying the Wuhan, D614G, Alpha and Beta spike, expressing the luciferase reporter gene. Luciferase was measured at 48h post infection. Data is mean ±SD of 2 independent experiments in triplicates. Statistical significance between the variants was evaluated by non-parametric Kruskal-Wallis test (ns, not statistically significant).

**Supplementary Table 1.**
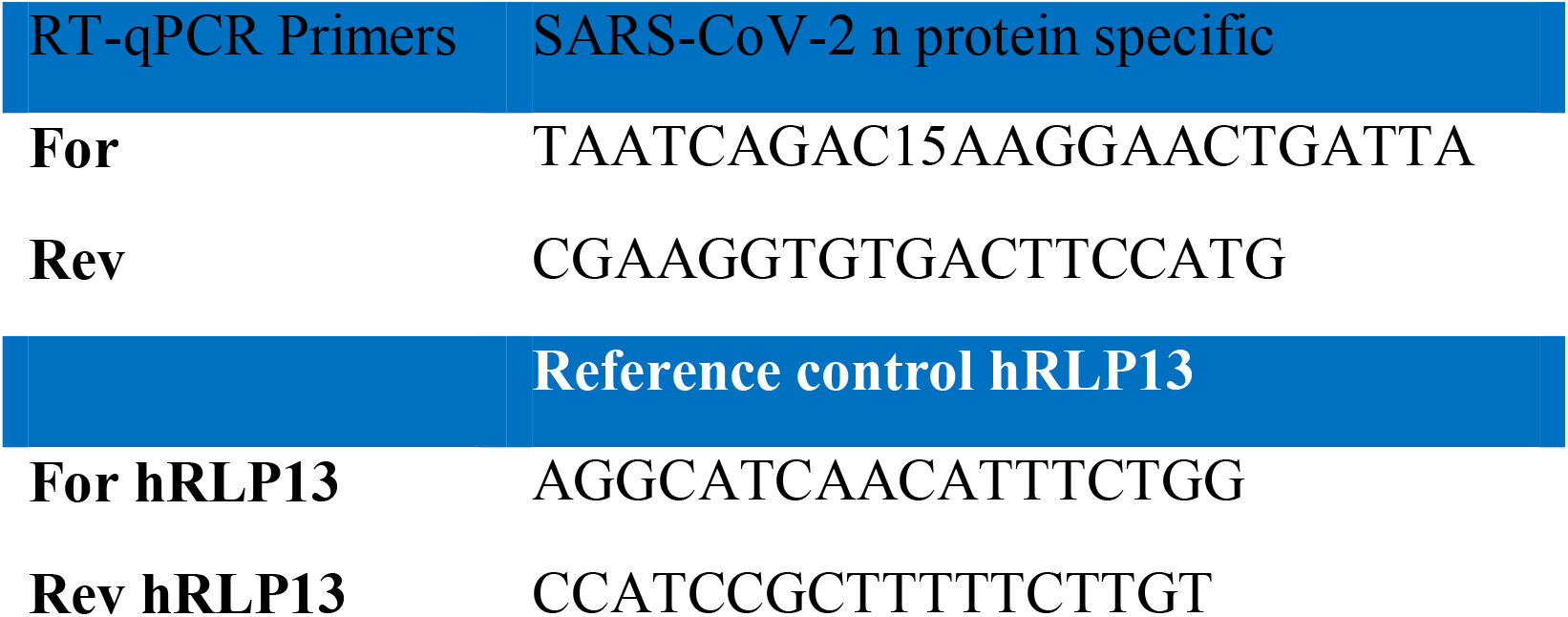
RT-qPCR oligomers.

## Notes

### Competing Interest Statement

The authors have declared no competing interest.

